# HIV-1 control in vivo is related to the number but not the fraction of infected cells with viral unspliced RNA

**DOI:** 10.1101/2024.07.01.601579

**Authors:** Adam A. Capoferri, Ann Wiegand, Feiyu Hong, Jana L. Jacobs, Jonathan Spindler, Andrew Musick, Michael J. Bale, Wei Shao, Michele D. Sobolewski, Anthony R. Cillo, Brian T. Luke, Christine M. Fennessey, Robert J. Gorelick, Rebecca Hoh, Elias K. Halvas, Steven G. Deeks, John M. Coffin, John W. Mellors, Mary F. Kearney

## Abstract

In the absence of antiretroviral therapy (ART), a subset of individuals, termed HIV controllers, have levels of plasma viremia that are orders of magnitude lower than non-controllers who are at higher risk for HIV disease progression. In addition to having fewer infected cells resulting in fewer cells with HIV RNA, it is possible that lower levels of plasma viremia in controllers is due to a lower fraction of the infected cells having HIV-1 unspliced RNA (HIV usRNA) compared with non-controllers. To directly test this possibility, we used sensitive and quantitative single cell sequencing methods to compare the fraction of infected cells that contain one or more copies of HIV usRNA in peripheral blood mononuclear cells (PBMC) obtained from controllers and non-controllers. The fraction of infected cells containing HIV usRNA did not differ between the two groups. Rather, the levels of viremia were strongly associated with the total number of infected cells that had HIV usRNA, as reported by others, with controllers having 34-fold fewer infected cells per million PBMC. These results reveal for the first time that viremic control is not associated with a lower fraction of proviruses expressing HIV usRNA, unlike what is reported for elite controllers, but is only related to having fewer infected cells overall, maybe reflecting greater immune clearance of infected cells. Our findings show that proviral silencing is not a key mechanism for viremic control and will help to refine strategies towards achieving HIV remission without ART.

## Significance statement

It is not fully understood what contributes to varying levels of viremia in untreated people with HIV (PWH). We investigated a possible association between high levels of plasma viremia and a high fraction of infected cells with proviruses expressing HIV-1 unspliced RNA (HIV usRNA), a requirement for virion packaging. We found that plasma viremia was not associated with the fraction of infected cells with HIV usRNA; but rather, only to the total number of infected cells. These findings refine our understanding of HIV-1 pathogenesis, suggesting that control occurs downstream of HIV usRNA synthesis.

## Main Text Introduction

Human immunodeficiency virus type 1 (HIV-1) primarily infects CD4+ T cells by binding to the CD4 receptor and a CCR5 or CXCR4 co-receptor, followed by fusion with the host cell membrane. The released viral core is then transported to the nucleus where reverse transcription is completed. After reverse transcription, the DNA form of the viral genome is integrated into host DNA to form a provirus. Host cell machinery is used for proviral transcription producing both spliced and unspliced forms of HIV-1 RNA. Unspliced RNA (usRNA) is packaged into virions, which can go on to infect new cells in the next round of viral replication. The steady-state level of plasma viremia is determined by the number of infected cells and the number of virions produced in each round of viral replication and is an important predictor of disease progression to acquired immunodeficiency syndrome (AIDS) and death (1, 2). In chronic untreated HIV-1 infection, the levels of plasma viremia can vary greatly from <50 to >10^6^ copies HIV RNA/mL across individuals (1, 2).

It has been shown that untreated people with HIV-1 (PWH) who have low levels of plasma viremia (viremic controllers, VC) have, on average, about 10-fold fewer infected cells per million CD4+ T cells than people with high levels of plasma viremia (non-controllers, NC) (3), and therefore, fewer CD4+ T cells with intracellular HIV RNA (4-6). We asked if, in addition to having fewer infected cells, VC have a smaller fraction of infected cells with detectable levels of HIV-1 unspliced RNA (HIV usRNA) compared to NC, suggesting that they may have a more effective block to proviral expression, as reported for elite controllers (7). Here, we sought to understand the relationship between the fraction of infected cells with HIV usRNA and the levels of plasma viremia in people with chronic untreated HIV-1 infection. To determine the fraction of infected cells with HIV usRNA, we used a sensitive, single infected cell sequencing method that we previously described (8). We focused on HIV usRNA (late transcript products) since this is the form that is translated into most virion proteins, packaged into virions, and detected as plasma viremia, while fully spliced RNA transcripts (early transcript products) may be present without resulting in virus production.

## Results

### Participant and sample characteristics

To determine the association between the levels of viremia and the fraction of infected cells with HIV usRNA, we assayed PBMC from 8 chronically infected, untreated donors with low levels of viremia (viremic controllers, VC, previously defined as having 50 to 2,000 copies HIV RNA/mL (9)), and 7 chronically infected, untreated donors with high levels of viremia (non-controllers, NC, >2,000 copies/mL) (**Table 1**). Although we assayed total PBMC, the vast majority of the infected cells in the peripheral blood are CD4+ T cells (10) and, therefore, the HIV usRNA detected can be considered to be from expression in this cell type. Direct assessment of PBMC avoids the cell losses that result from enrichment or cell sorting protocols. Results were normalized for the number and percent of CD4+ T cells, where indicated in each analysis and figure below.

**Table 1.** Participant characteristics.

| Group* | Participant Identifier (PID) | Cohort Location | Age (years) | Sex<sup>†</sup> | Race<sup>‡</sup> | Minimum duration of infection (years) | CD4+ T cells at sampling (cells/μL) | %CD4+ T cells at sampling | HIV plasma RNA copies/mL at sampling<sup>§</sup> | HLA-B alleles<sup>¶</sup> |
| --- | --- | --- | --- | --- | --- | --- | --- | --- | --- | --- |
| Viremic controllers (VC) | 1508 | SCOPE, UCSF | 53 | M | Black | 30 | 484 | 19 | 128 | B*58/B*81 |
|  | 1517 | SCOPE, UCSF | 47 | M | White | 25 | 946 | 44 | 60 | B*18/B*57 |
|  | 1775 | SCOPE, UCSF | 37 | M | White | 5 | 1,066 | 41 | 1,704 | B*14/B*57 |
|  | 1788 | SCOPE, UCSF | 51 | M | White | 16 | 1,136 | 39 | 331 | B*27/B*57 |
|  | 3713 | SCOPE, UCSF | 28 | M | White | 1 | 762 | 29 | 122 | B*27/B*27 |
|  | 3714 | SCOPE, UCSF | 60 | M | White | 4 | 1,008 | 40 | 854 | ND |
|  | 515 | University of Pittsburgh | 28 | M | White | 2 | 749 | 30 | 1,951 | ND |
|  | 826 | University of Pittsburgh | 54 | M | Black | 18 | 1,053 | 44 | 120 | ND |
| **Median** |  |  | **49** |  |  | **11** | **977** | **40** | **230** |  |
| **[IQR]** |  |  | **[35-53]** |  |  | **[4-20]** | **[759-1,056]** | **[30-42]** | **[122-1,067]** |  |
| Non-controllers (NC) | 291 | University of Pittsburgh | 53 | M | Black | 9 | 874 | 28 | 184,243 | ND |
|  | 416 | University of Pittsburgh | 44 | M | Black | 3 | 870 | 22 | 13,164 | ND |
|  | 801 | University of Pittsburgh | 50 | M | Black | 28 | 256 | 25 | 5,727 | ND |
|  | 740 | University of Pittsburgh | 51 | M | Black | 30 | 214 | 10 | 32,329 | ND |
|  | 477 | University of Pittsburgh | 26 | F | Black | 5 | 114 | 9 | 139,845 | ND |
|  | 3611 | SCOPE, UCSF | 37 | M | White | 4 | 118 | 7 | 275,685 | B*08/B*44 |
|  | 1226 | SCOPE, UCSF | 38 | M | Multiracial | 12 | 1,155 | 53 | 165,418 | B*15/B*44 |
| **Median** |  |  | **44** |  |  | **9** | **256** | **22** | **139,845** |  |
| **[IQR]** |  |  | **[38-51]** |  |  | **[5-20]** | **[166-872]** | **[10-27]** | **[22,747-174,831]** |  |

The donor characteristics are consistent with the time of sampling for **Table 2**.
\* Groups include viremic controllers (plasma RNA 60-2,000 copies/mL not on ART) and non-controllers (plasma RNA >5,000 copies/mL not on ART).
<sup>†</sup> Sex assigned at birth: M (male), F (female).
<sup>‡</sup> Race as reported by donors.
<sup>§</sup> Level of plasma viremia was determined by either Abbott® Real-time HIV-1 Assay (SCOPE, UCSF) or COBAS® HIV-1 Test (University of Pittsburgh).
<sup>¶</sup> ND (not determined).

Levels of viremia in VC were low but detectable (median 230, range 60-1,951 copies/mL) and levels of viremia in NC were high (median of 139,845 copies/mL, range 5,727-275,685 copies/mL). Participants were mostly assigned male at birth (n=14/15). Race as reported by donors was black (n=7/15), white (n=7/15), and multiracial (n=1/15). Donors were diagnosed with HIV-1 subtype B with a minimum duration of infection median of 9 years [IQR 4-22 years] prior to sample collection. Donors were either ART naïve or not currently on an ART regimen due to a planned or unplanned ART interruption. One donor (PID 801) had previous virologic failures and developed multiple drug resistance mutations but had been off ART for 3 years at the time of sampling.

### Virologic characterization of VC and NC

Prior to comparing the fraction of infected cells with HIV usRNA in the VC vs. NC, we measured the levels of plasma viremia in both groups as HIV RNA copies per mL by either the Abbott® real-time assay or COBAS® HIV-1 test (**Fig. 1A**) and estimated the number of infected cells (provirus+ cells) by measuring HIV integrase DNA copies per million PBMC by digital PCR (**Fig. 1B**) (11). We used HIV DNA levels as a surrogate for the number of infected cells based on the finding that most infected cells have only a single HIV DNA copy even prior to ART (10). Plasma viremia was significantly lower in VC compared to NC (p=0.0003, Mann-Whitney). The number of infected cells was also significantly lower in VC (median 42 HIV-1 DNA/10^6^ PBMC) compared to NC (median 1,140 HIV-1 DNA/10^6^ PBMC) (p=0.0003, Mann-Whitney). We found no association between the number of infected cells and the minimum duration of infection (Spearman’s ρ=0.11, p=0.68; **Fig. 1C**). However, we did find a trend towards a difference in the CD4+ T cell count and the %CD4+ T cells between controllers and non-controllers (**Fig. S1**). As expected from previous studies (12, 13), there was a significant association between the level of plasma viremia and the number of infected cells (Spearman’s ρ=0.84, p=0.0002; **Fig. 1D**). However, this finding does not differentiate among infected cells as to their transcriptional activity or presumed virion production.

**Figure 1.**
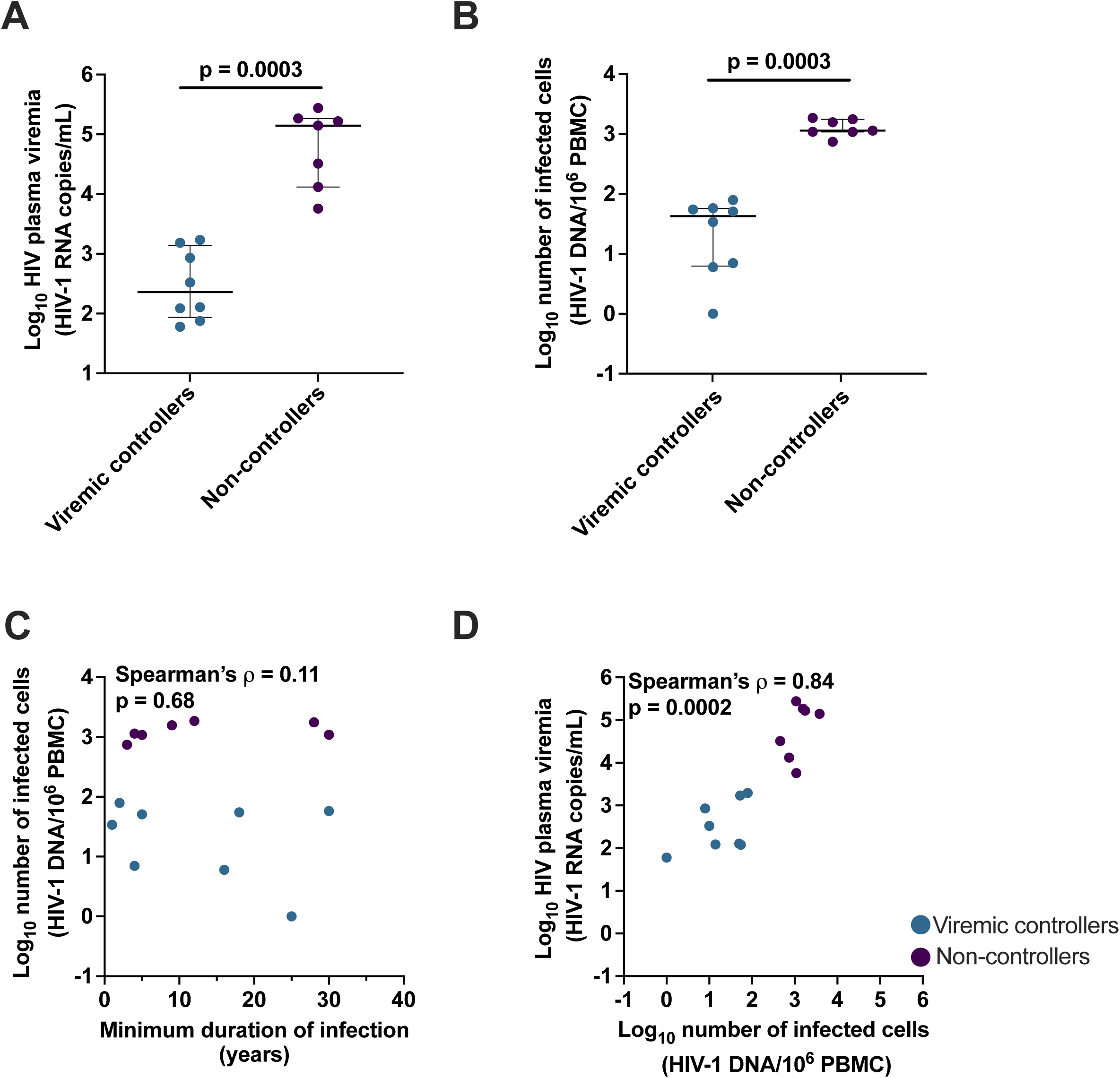
Participant baseline virologic characteristics. (**A**) Level of plasma viremia in HIV-1 RNA copies per mL, with p-value (Mann-Whitney test) shown above. Median and interquartile range are also shown. (**B**) Number of provirus+ cells (a surrogate for the number of infected cells) by HIV-1 DNA per million PBMC, Mann-Whitney test with median and interquartile range is shown. (**C**) Minimum duration of HIV infection (years) and number of provirus+ cells by HIV-1 DNA per million PBMC. (**D**) Number of provirus+ cells by HIV-1 DNA per million PBMC and level of plasma viremia by HIV-1 RNA copies/mL. Each symbol represents an individual donor within the respective group.

### The total number of PBMC with HIV usRNA is lower in VC than in NC and is correlated with levels of viremia

Before measuring the fraction of infected PBMC with HIV usRNA, we asked if the total number of PBMC with HIV usRNA correlated with their different levels of viremia using our CARD-SGS assay (8), as reported using RNA-flow FISH approaches (4-6). Detailed methods for the CARD-SGS assay are included in the methods section but, in brief, PBMC are spread across 8-10 aliquots with each aliquot containing only a single or a few infected cells with HIV usRNA. The total nucleic acid is extracted from each aliquot, the DNA is digested, and cDNA is synthesized using a gene-specific primer in the *pol* gene. All HIV cDNA molecules are then counted and sequenced using single-genome sequencing (SGS) (8).

Consistent with others (4-6), we found that the total number of PBMC with HIV usRNA molecules was significantly lower in VC (median 2.6 cells with HIV usRNA /10^6^ PBMC) compared to NC (median 89.2 cells with HIV usRNA /10^6^ PBMC) (p=0.0003, Mann-Whitney; **Fig. 2A**). Consequently, we found a strong positive association between levels of plasma viremia and the total number of cells with HIV usRNA (Spearman’s ρ=0.77, p=0.001; **Fig. 2B**), although the correlation was slightly less positive than with the total number of infected cells reported above. As expected, we also found a correlation between the total number of infected cells and the number that contained HIV usRNA (Spearman’s ρ=0.92, p<0.0001; **Fig. 2C**). After normalizing for the %CD4+ T cells, we also found that the number of HIV usRNA copies/10^6^ CD4+ T cells was significantly lower in VC (median 0.9 HIV usRNA/million CD4+ T cells) compared to NC (19.1 usRNA/million CD4+ T cells) (p=0.001, Mann-Whitney; **Fig. S2A**).

**Figure 2.**
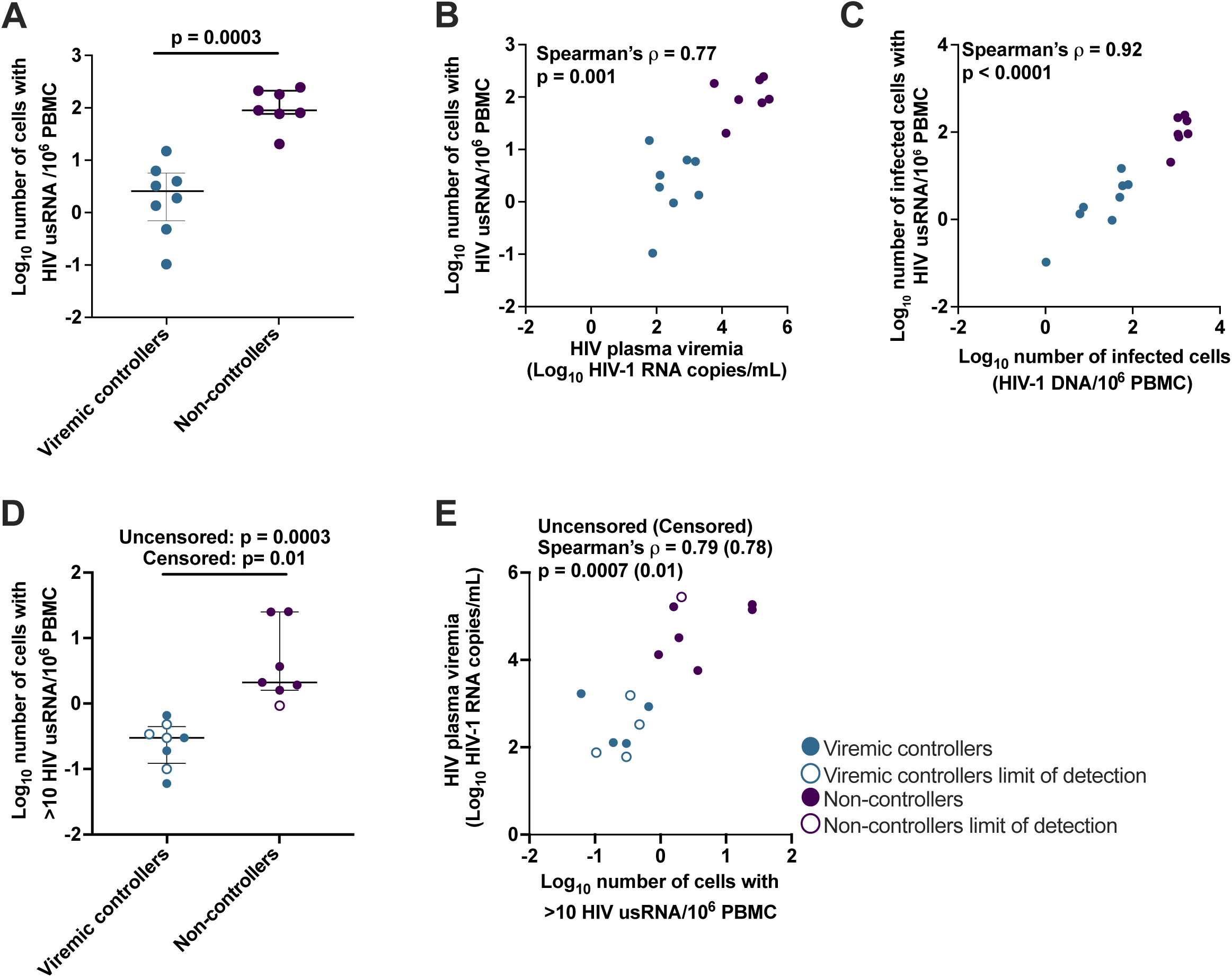
Number of PBMC with HIV usRNA and association with levels of plasma viremia. (**A)** Number of cells with HIV usRNA per million PBMC, Mann-Whitney test, median and interquartile range are shown. (**B**) The log10-transformed number of cells with HIV usRNA per million PBMC and log10-transformed plasma viremia as HIV-1 RNA copies per mL in VC and NC. (**C**) Provirus+ cells by HIV-1 DNA per million PBMC and log10-transformed number of provirus+ cells with HIV usRNA per million PBMC in VC and NC. (**D**) Number of cells per million PBMC that are “high-expressing” (>10 HIV usRNA), Mann-Whitney test with median and interquartile range are reported. (**E**) Number of “high-expressing” cells per million PBMC and log10-transformed plasma viremia in HIV-1 RNA copies per mL of plasma. Open shapes indicate no detectable “high-expressing” cells in a particular donor; thus, a lower limit of detection estimate (determined as 1/number of PBMC assayed) is shown. A Mann-Whitney test was performed for both uncensored and censored data (excluding no undetectable “high-expressing” cells donors). Each symbol represents an individual donor within the respective group.

Additionally, we investigated the frequency of cells with >10 HIV usRNA by assuming that identical HIV usRNA sequences within the same aliquot of PBMC originated from the same cell, made possible because of the high HIV genetic diversity in these chronically untreated individuals (more on the HIV genetic diversity below). We termed these cells “high-expressing”. We found significantly fewer “high-expressing” cells per million PBMC in VC (median 0.3 “high-expressing” proviruses/10^6^ PBMC) compared to NC (median 2.1 “high-expressing” proviruses/10^6^ PBMC) (p=0.0003, Mann-Whitney; **Fig. 2D**). After normalizing for %CD4, we also found significantly fewer “high-expressing” cells/million CD4+ T cells in VC (median 1.3 “high-expressing” proviruses/10^6^ CD4+ T cells) compared to NC (median 8.3 “high-expressing” proviruses/10^6^ CD4+ T cells) (p=0.0007, Mann-Whitney; **Fig. S2B**). The number of “high-expressing” cells/million PBMC was positively associated with levels of plasma viremia (Spearman’s ρ=0.79, p=0.0007; **Fig. 2E**) which held true after normalizing for %CD4 (Spearman’s ρ=0.60, p=0.02). Our results are consistent with lower measurements of intracellular HIV usRNA in bulk cell populations in VC compared to NC (11, 14-16) and with results reported using RNA-flow FISH approaches in untreated PWH (4-6).

### The fraction of infected PBMC with HIV usRNA is not different in VC and NC

Since previous reports suggest that elite controllers have a block to proviral expression related to their different profile of integration sites (7), we next measured the fraction of infected PBMC with HIV usRNA in the VC vs. NC to determine if VC have a similar block to proviral expression as reported for the elite controllers. If so, we would expect that the VC would have a smaller fraction of cells with HIV usRNA compared to the NC. We determined the fraction of infected cells that contained HIV usRNA by using the estimated number of infected cells in the aliquots as the denominator and the number of unique RNA sequences in each aliquot as the numerator (8, 11). Because unique RNA sequences are considered to originate from different infected cells (most cells containing a different provirus in untreated, chronic infection (10)), we restricted our study to donors with high levels of HIV genetic diversity as stated above. It has been shown that many proviruses in untreated infection are intact (17-19) and that some defective proviruses can express HIV usRNA (17, 20-22), therefore, we did not distinguish between HIV usRNA from intact vs. defective proviruses in this study. Because of the very low number of infected cells in the VC (some with <10 infected cells per million PBMC) and because of the rarity of accessing samples from donors in chronic HIV infection who are not on ART, we assayed as many PBMC as obtainable from these historic samples (3-16 million). However, our depth of sampling of HIV usRNA+ cells in VC compared to the NC was still a median of ∼4-fold lower (**Table 2**). To account for the lower level of sampling in the VC, we normalized for the number of infected cells with HIV usRNA across the two groups.

**Table 2.** Fraction of infected cells with HIV usRNA in viremic controllers and non-controllers.

| Group | Participant Identifier (PID) | Total number of cells assayed | Estimated number of infected cells assayed | Number of HIV usRNA sequences obtained | Estimated number of infected cells with HIV usRNA * | Estimated % of infected cells with HIV usRNA | Average number of HIV usRNA copies per cell [range] † | Median number of HIV usRNA copies per cell [IQR] † |
| --- | --- | --- | --- | --- | --- | --- | --- | --- |
| Viremic controllers (VC) | 1508 | 2,900,000 | 248 | 31 | 17 | 7 | 1.7 [1-6] | 1 [1-1.75] |
|  | 1517 | 9,644,140 | 10 | 1 | 1 | 10 | 1.0 [1-1] | 1 [1-1] |
|  | 1775 | 5,257,870 | 268 | 40 | 17 | 6 | 2.4 [1-11] | 1 [1-1] |
|  | 1788 | 16,194,930 | 97 | 46 | 22 | 23 | 2.1 [1-24] | 1 [1-1] |
|  | 3713 | 2,075,645 | 71 | 2 | 2 | 3 | 1.0 [1-1] | 1 [1-1] |
|  | 3714 | 10,043,290 | 70 | 80 | 19 | 27 | 4.2 [1-17] | 1 [1-5] |
|  | 515 | 3,040,000 | 240 | 90 | 19 | 8 | 4.7 [1-38] | 1 [1-3] |
|  | 826 | 3,280,000 | 180 | 84 | 49 | 27 | 1.7 [1-7] | 1 [1-2] |
|  | Median | 4,268,935 | 139 | 43 | 18 | 9 | 2 |  |
| Non-controllers (NC) | 291 | 158,400 | 248 | 189 | 39 | 16 | 4.8 [1-48] | 1 [1-1] |
|  | 416 | 2,160,000 | 1,604 | 156 | 44 | 3 | 3.5 [1-62] | 1 [1-2] |
|  | 801 | 544,000 | 594 | 206 | 98 | 16 | 3.1 [1-23] | 1 [1-2] |
|  | 740 | 1,040,000 | 475 | 189 | 93 | 20 | 2.0 [1-30] | 1 [1-1] |
|  | 477 | 80,000 | 306 | 139 | 17 | 6 | 8.2 [1-77] | 1 [1-1] |
|  | 3611 | 1,252,414 | 1,427 | 139 | 79 | 7 | 1.8 [1-20] | 1 [1-1] |
|  | 1226 | 473,733 | 877 | 43 | 40 | 5 | 1.1 [1-2] | 1 [1-1] |
|  | Median | 544,000 | 594 | 156 | 44 | 7 | 3 |  |

\* Assuming that identical sequence are from a single cell.
† Cells without HIV usRNA were excluded.

We found that only a small fraction of the infected PBMC contained HIV usRNA in both the VC and the NC, and that these fractions were not significantly different between the groups: VC – median 9% IQR [6-26%]; NC - median 7% IQR [5-16%] (p=0.41, Mann-Whitney; **Table 2 and Fig. 3A**). These results indicate that levels of plasma viremia are not related to the fraction of infected PBMC that have HIV usRNA (Spearman’s ρ= -0.32, p=0.24; **Fig. 3B**). They also indicate that most infected PBMC in chronic untreated infection contain proviruses that are not expressing HIV usRNA at the time of sampling, independent of plasma viremia levels, consistent with reports by others (4-6). These results refute our idea that VC have a higher fraction of latent proviruses than NC and demonstrate that, on average, >90% of infected PBMC in both groups have proviruses that are not expressing HIV usRNA at a given point in time, even in the absence of ART.

**Figure 3.**
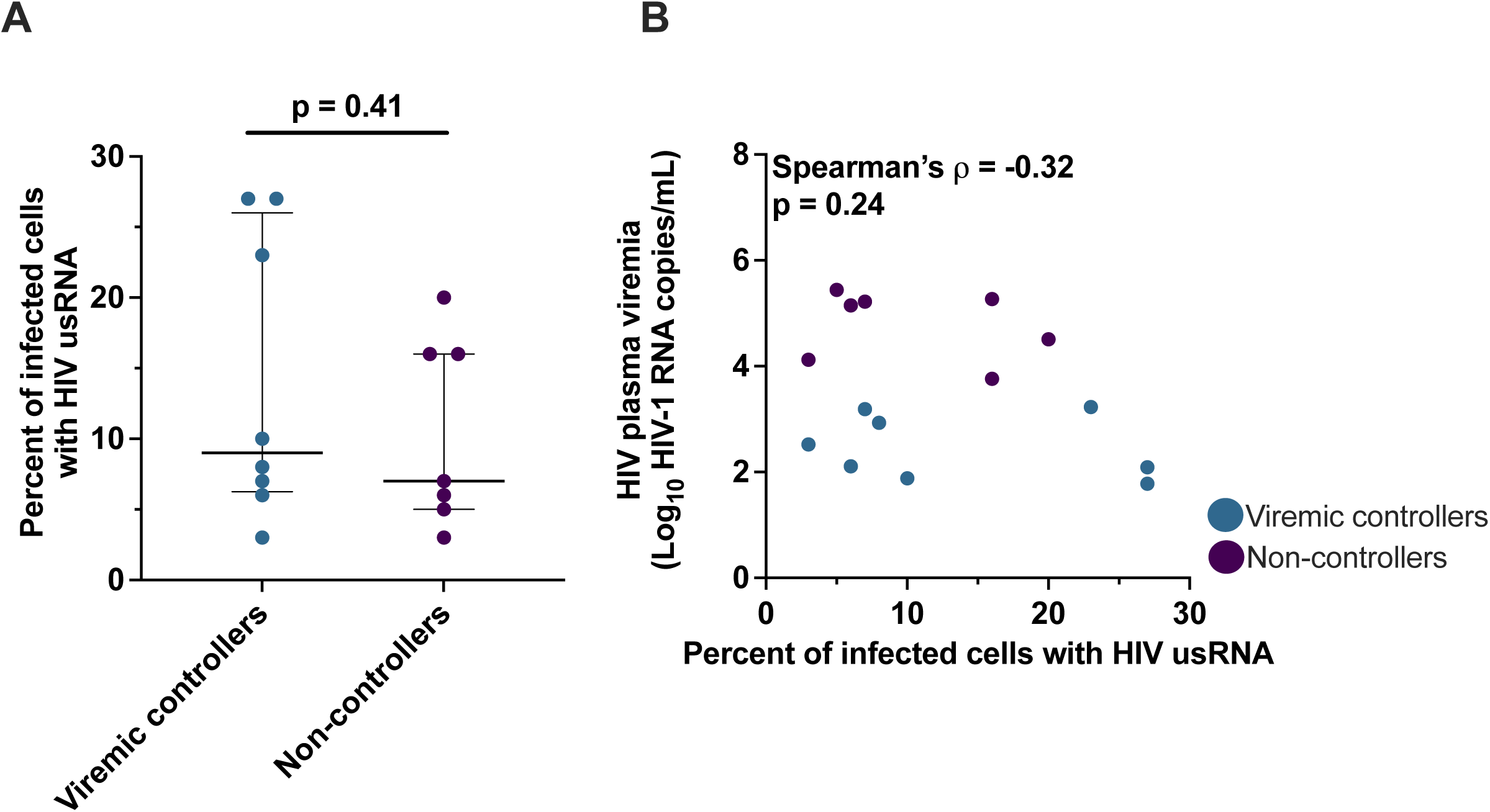
The fraction of infected cells with HIV usRNA is not associated with levels of plasma viremia. **(A)** Percent of infected cells (provirus+ cells) with HIV usRNA in VC and NC. Median and interquartile range are shown. **(B)** Percent infected cells with usRNA and the log10-transformed HIV plasma viremia as HIV-1 RNA copies per mL in VC and NC. Each symbol represents an individual donor within the respective group.

### The levels of HIV usRNA in single infected cells are not different in VC and NC

By examining the identical RNA sequences from the results of the CARD-SGS assay, the number of HIV usRNA molecules could be estimated for each infected cell (**Fig. 4A**). The average number of HIV usRNA copies per HIV usRNA+ cell was not significantly different between VC (2 copies/cell) and NC (3 copies/cell) (p=0.27, Mann-Whitney; **Table 2 and Fig. 4B**). Of 147 infected PBMC with HIV usRNA in VC and 410 in NC, the frequency of “high-expressing” cells (defined here as >10 HIV usRNA molecules/cell) was 4.8% and 3.5% respectively (p=0.46) (**Fig. 4C**). Despite not having significantly different levels of HIV usRNA in the vast majority of single cells, there was a trend towards the “high-expressing” cells having fewer copies of HIV usRNA in VC (mean 19.6 HIV usRNA copies/cell) vs. NC (mean 33.5 HIV usRNA copies/cell) (p=0.06, chi-square test with 1 degree of freedom), although this trend did not reach significance. These results show that varying levels of plasma viremia in VC vs. NC are not significantly associated with the level of HIV usRNA expression in single infected cells.

**Figure 4.**
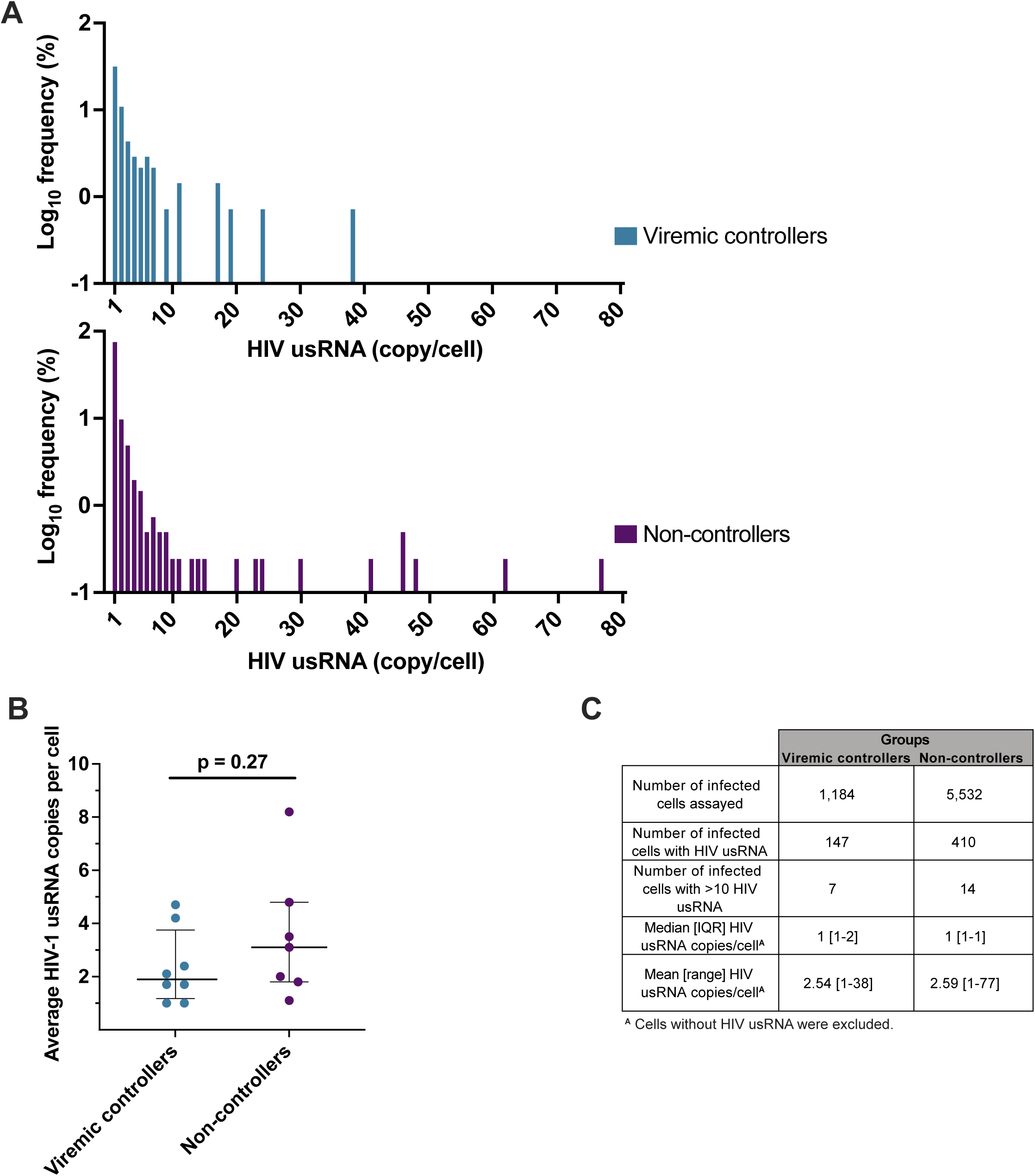
Levels of HIV usRNA in single cells. (**A**) Distribution plots of the frequency of HIV usRNA copies per cell in VC and NC. (**B**) Average number of HIV-1 usRNA copies per cell detected for each participant. Mann-Whitney test with median and interquartile range are shown. (**C**) Comparison of HIV usRNA in single cells between VC and NC. The number of infected cells assayed with the indicated number expressing HIV usRNA is reported. The number of “high-expressing” cells (>10 copies HIV usRNA/cell) in each group with the median and interquartile range, mean and range for the HIV usRNA copies per cell are reported. Each symbol represents an individual donor within the respective group.

### Phylogenetic analysis of HIV usRNA reveals expression from a diverse population of proviruses

To investigate the genetic diversity of HIV usRNA in VC and NC, we sequenced and phylogenetically analyzed plasma HIV RNA (red circles), PBMC HIV DNA (black triangles), and intracellular HIV usRNA (different colored rectangles representing different aliquots of cells) in VC (**Fig. 5**) and NC (**Fig. 6**). Sampling in VC was shallow compared to NC due to the low number of infected cells in these individuals. We performed SGS on multiple 96-well PCR plates at the endpoint for RNA and DNA (yielding ∼6—100 sequences per sample) to measure the average pairwise distance (APD). For HIV usRNA, we performed SGS on 8-10 aliquots of PBMC, each containing, on average, <5 cells with HIV usRNA. In total, we aimed to sequence about 20 genomes from each compartment to determine if there was a difference in the genetic diversity between intracellular HIV usRNA, proviral DNA, and/or plasma RNA. However, in a few samples, few or no sequences were obtained due to low viral loads and/or limited sample availability. APDs were measured where >10 sequences per compartment were obtained. Identical HIV usRNA and plasma sequences were collapsed prior to calculating APD so that multiple transcripts from the same cell were not overrepresented.

**Figure 5.**
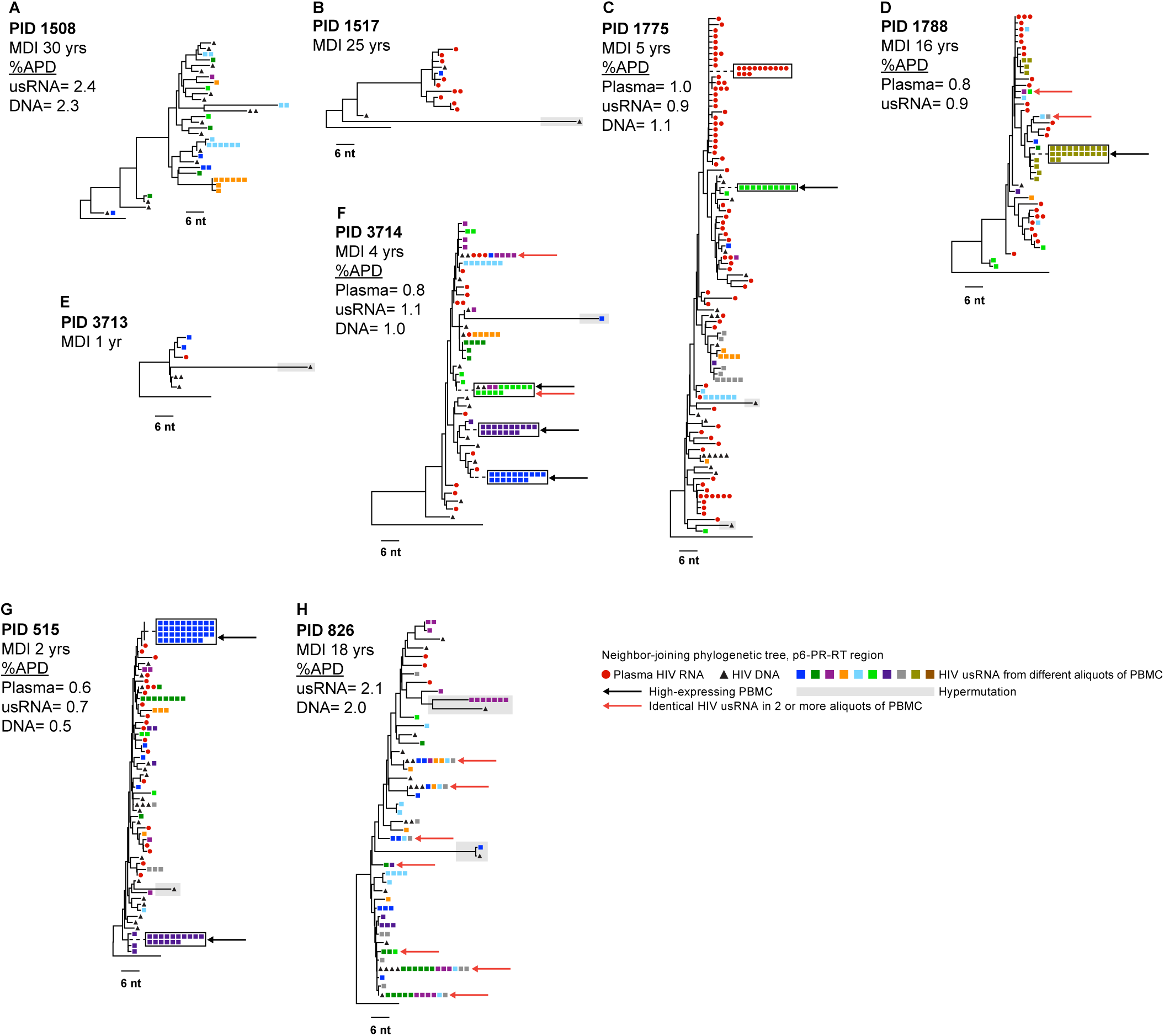
Distance trees of viremic controllers. Neighbor-joining p-distance tree of the p6-PR-RT region of HIV-1 obtained from CARD-SGS assay on the viremic controllers. The Participant ID (PID) is denoted with the minimum duration of infection (MDI) in years. The black triangles show the proviruses from PBMC, red circles are the plasma virus, and the squares are the intracellular HIV usRNA from PBMC with each color square representing a different aliquot. Black arrows indicate “high-expressing” PBMC and red arrows indicate identical HIV usRNA sequences found in >1 aliquot of PBMC.Diversity was measured where appropriate (see Materials and Methods) by the average pairwise distance with predicted hypermutants removed and identical sequences collapsed.

**Figure 6.**
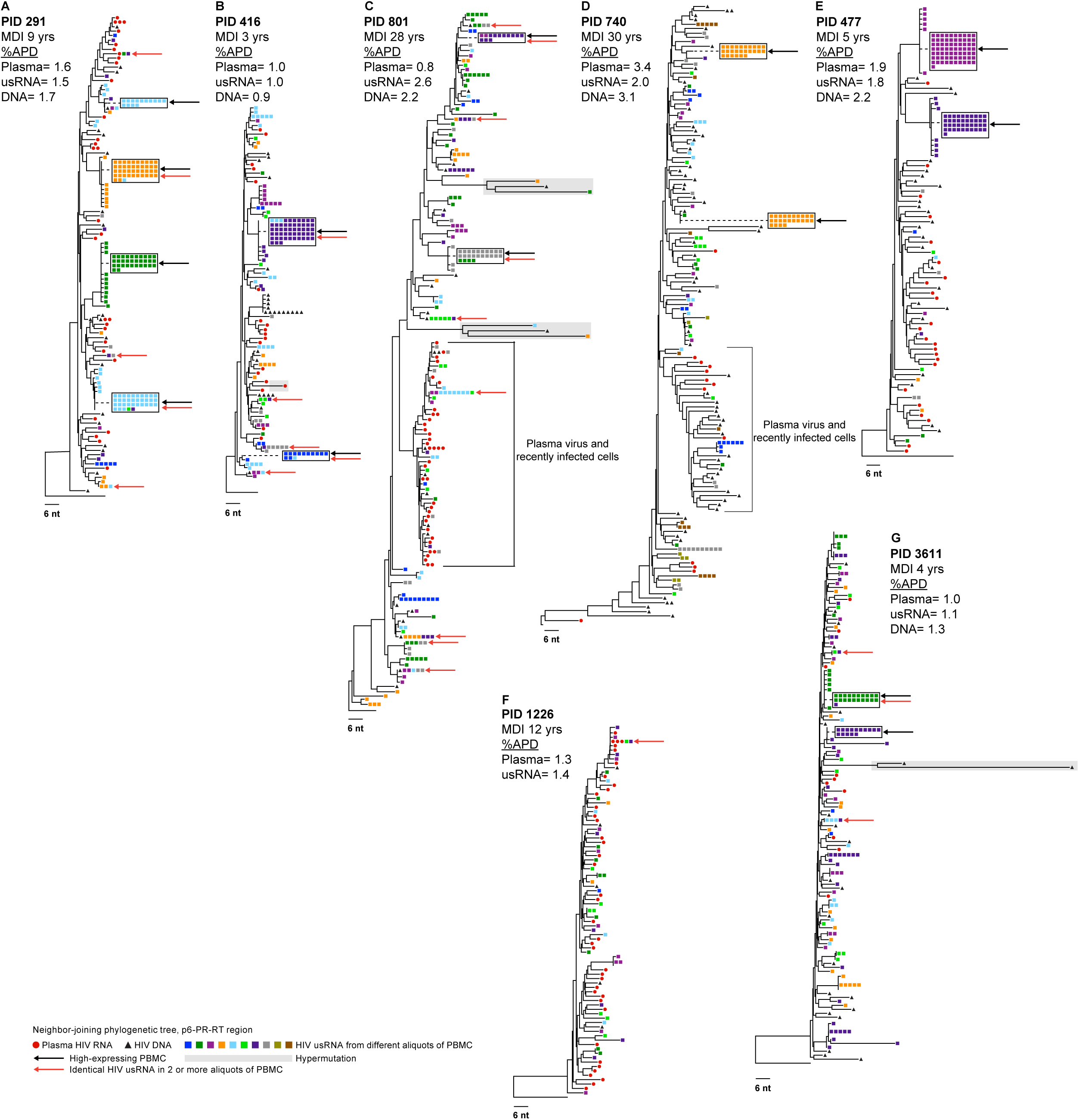
Distance trees of non-controllers. Neighbor-joining p-distance tree of the p6-PR-RT region of HIV-1 obtained from CARD-SGS assay on the non-controllers. The Participant ID (PID) is denoted with the minimum duration of infection (MDI) in years. The black triangles show the proviruses from PBMC, red circles are the plasma virus, and the squares are the intracellular HIV usRNA from PBMC with each color square representing a different aliquot. Black arrows indicate “high-expressing” PBMC and red arrows indicate identical HIV usRNA sequences found in >1 aliquot of PBMC. Diversity was measured where appropriate (see Materials and Methods) by the average pairwise distance with predicted hypermutants removed and identical sequences collapsed.

Overall, the similar %APDs of single-genome sequences in plasma HIV RNA, HIV DNA, and intracellular HIV usRNA within donors indicates that diverse populations of proviruses were transcriptionally active in both VC and NC (**Fig. 5 & 6**). The APDs of the HIV usRNA were not significantly different, ranging from 0.7 to 2.4% in the VC and from 1.0% to 2.6% in the NC (p=0.31, Mann-Whitney), typical for *pol* diversity in chronic infection. In trees where all compartments were adequately sampled, HIV usRNA appears to have originated from both recently infected cells and archival proviruses. Two specific examples of what appear to be archival infected cells are seen in PID 801 (**Fig. 6C**) and PID 740 (**Fig. 6D**). In these two individuals, most or all plasma sequences formed well-defined nodes. The absence of corresponding plasma virus sequences elsewhere in these two trees implies that HIV usRNA and DNA on other nodes may be from cells infected at an earlier timepoint. These findings suggest that the ∼10% of infected cells that have HIV usRNA comprise a diverse population of infected cells, both recently infected and archival cells that are long lived and/or clonally expanded. Evidence that these cells may be clonally expanded is found where identical usRNA sequences are present across multiple aliquots of infected cells (different colored squares). Such cases are indicated with red arrows and are found in both the VC and NC groups. This finding suggests that clones of cells possibly infected at an earlier time persist and express HIV usRNA in untreated infection, as has been shown to occur during ART (8).

HIV usRNA from “high-expressing” cells (indicated with black arrows) were found in both VC (n=4/5 donors with adequate sampling) and NC (n=6/7 donors) and, on occasion, had sequences matching RNA in other cells, implying either clonal spread of virus or expansion of infected cells. Because “high-expressing” cells are almost never detected in people with their viremia suppressed on ART (8), these cells may be in the late stage of the viral replication cycle and rapidly die by cytopathicity of viral replication or after being recognized and killed by immune cells .

## Discussion

While the number of total CD4+ T cells with HIV RNA had previously been quantified and shown to be associated with levels of viremia (4-6), the fraction of infected cells with HIV usRNA and the levels of usRNA in individual infected cells from viremic individuals had not been carefully quantified until the current work. We found that, on average, only about 10% of infected PBMC from individuals with chronic viremia who were not receiving ART had detectable levels of HIV usRNA, similar to prior studies of donors on long-term ART (23-25). The proportion of infected cells expressing HIV usRNA was independent of levels of plasma viremia over a range of more than 4,000-fold, from 60 to 275,000 HIV-1 RNA copies/mL. We had expected that, if there was a further block to proviral expression in VC compared to NC, as reported for elite controllers (7), we would find a smaller fraction of infected cells with detectable levels of HIV usRNA, but no such association was found. Instead, we found that levels of detectable plasma viremia are directly correlated only with the total number of infected PBMC (consistent with findings of others (12, 13)) and the total number with HIV RNA (also consistent with others) (4-6). Significant correlations were also found between the levels of viremia and the total number of PBMC with >10 HIV usRNA molecules. These observations reveal that the lower levels of viremia in VC compared to NC are not from greater silencing of proviral expression of HIV usRNA, rather only due to fewer infected cells in VC compared to NC. Before the current work, the distinction between the two potential mechanisms explaining differences in viremia had not been made (**Fig. 7**).

**Figure 7.**
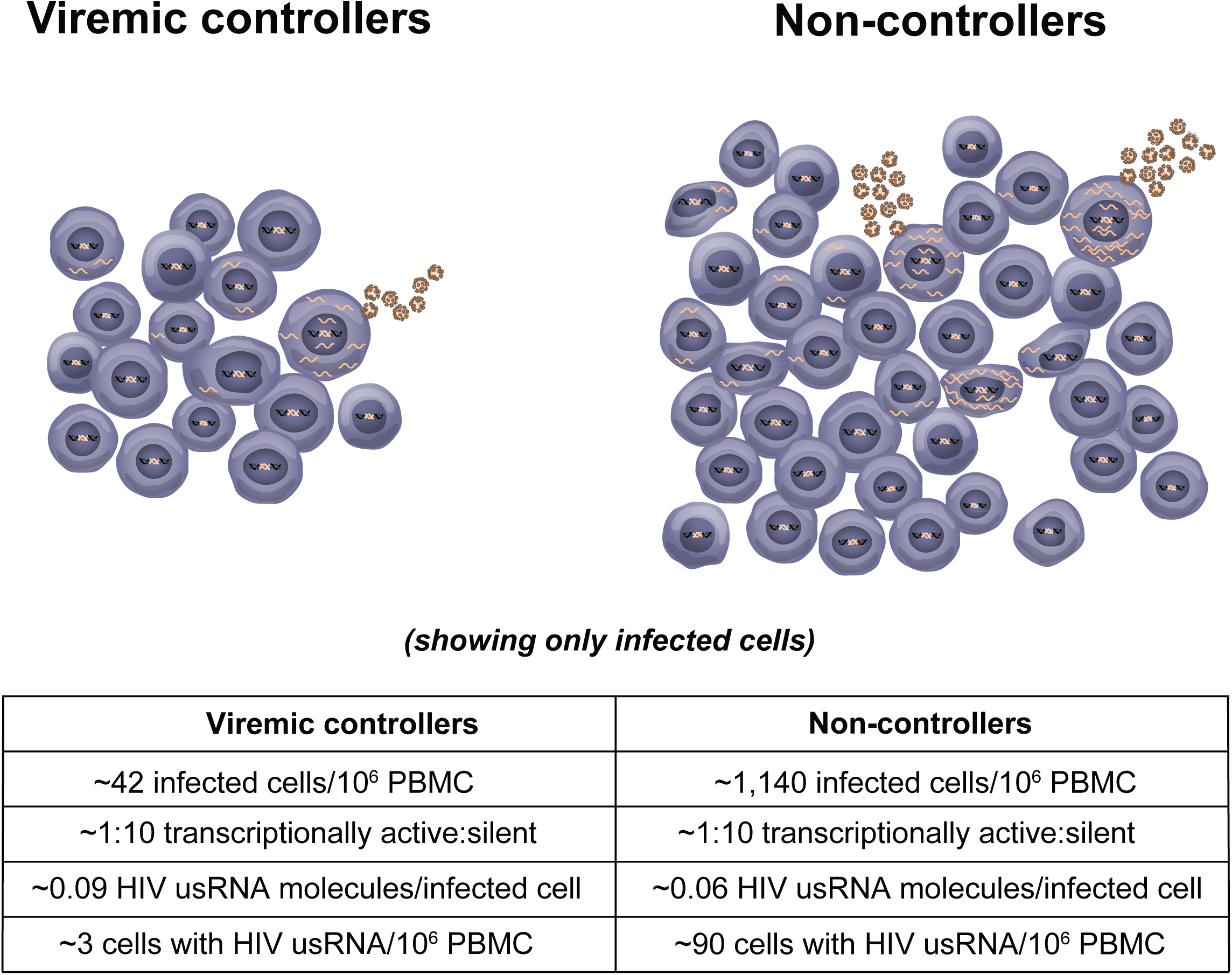
HIV-1 control is related to the number but not the fraction of infected cells with transcriptionally active proviruses. There are approximately 27-fold fewer infected cells in the viremic controllers. The ratio of transcriptionally active to silent proviruses is the same in the two groups with similar levels of HIV usRNA molecules per infected cell. These model based on measured at the time of sampling from Table 2.

We found that PBMC with higher levels of HIV usRNA (>10 HIV RNA molecules detected per cell) were rare, likely because they die rapidly, leaving only a small window of time in which they can be detected. We hypothesize that these “high-expressing” cells are in the late stages of the viral replication cycle while those with no detectable HIV usRNA are either early in the viral replication cycle or are long-lived, latently infected cells. However, it is also possible that the “high-expressing” cells are of a specific cell type or differentiation state that simply allow for higher levels of proviral expression (26). The high frequency of infected cells with low levels of HIV usRNA may reflect the steady-state levels of HIV usRNA expression, suggesting that these molecules may not always accumulate, but rather are encapsulated or degraded shortly after their synthesis. While cells with low levels of HIV usRNA are also frequently detected in individuals on ART, “high-expressing” cells have only rarely been detected in these individuals (8, 24, 25), an unsurprising result, given that the level of viremia in individuals on ART is, on average, about 10,000-fold lower than in the absence of ART.

In contrast to the situation on ART, the samples analyzed here were from donors with ongoing viral replication where about half of proviruses have been reported to be intact (17-19). However, it has been demonstrated that defective proviruses can accumulate prior to ART initiation (19) and that some defective proviruses can express HIV usRNA (17). Our methods distinguish between proviruses that express HIV usRNA and those that do not, but do not always distinguish between expression of defective and intact proviruses, or transcripts that aborted downstream from *pol*. Although at least half of the HIV usRNA that we detect is likely derived from intact proviruses, HIV usRNA from defective proviruses that retain functional *tat* and *rev*, as well as an intact major splice donor site, may also undergo proviral expression of HIV usRNA and may also contribute to plasma HIV RNA levels (27, 28).

Infected cells that are not positive for HIV usRNA may comprise several populations: 1) cells infected in the day(s) just prior to sampling, including those still undergoing viral DNA synthesis, integration, or incomplete transcription, 2) cells infected weeks, months, or years prior to sampling with integrated but silenced proviruses (some of which may be in cell clones), 3) cells with defective proviruses that cannot be transcribed, and 4) cells in which HIV DNA integration has failed leading to its circularization (29). In untreated PWH, the relative proportion of cells with defective or latent proviruses to recently infected cells and those with unintegrated linear or circular DNA will increase over time, due both to their longer lifetime (30), and to clonal expansion during untreated infection, as well as during ART (31).

The “high-expressing” cells seen in this study are either recently infected or were infected days, weeks, months, or years prior to sampling, underwent clonal expansion (32, 33), and chronically express or were recently reactivated to express HIV usRNA. The evidence of clonal expansion is in the form of identical RNA sequences in two or more aliquots of cells in both VC and NC (red arrows in **Fig. 5 & 6**). In two of the trees, the plasma RNA sequences, the large majority of which must have been derived from recently infected cells, are clustered separately from the high expressing cells, consistent with the high expressing cells possibly being infected at a different time. In PID 801 (**Fig. 6C**) who was diagnosed with HIV 28 years prior and subjected to multiple cycles of ART, the clustering of all the plasma RNA sequences into a single node was likely a result of evolutionary pressure related to the on and off ART regimen endured by this participant. The presence of the high-expressing cells detected in this participant as well as 8 of the clonally expanded HIV usRNA expressing cells falling outside of this node implies that only a small minority of the usRNA detected in this individual originated from recently infected cells. In PID 740, living with HIV for ≥30 years, the majority (8/12) of the plasma RNA sequences were also found in a node that lacked the “high-expressing” cells. These examples suggest that, in some cases, long-term infected cells can contribute significantly to the pool of cell-associated HIV usRNA, but much less so to the production of plasma virus. Further experimentation, such as longitudinal analysis by CARD-SGS following ART initiation, will be necessary to establish the generality of this observation.

Our previous study showed that the CARD-SGS assay can detect a single HIV RNA molecule in a single-infected cell (8). We used this approach in preference to single-cell RNA-seq since most cells interrogated by RNA-seq are uninfected (99.9% or more) (34-37); however, if many cells are sampled in RNA-seq datasets, HIV transcripts may be detected in those that are infected, though not with single RNA molecule sensitivity or (37-40). In contrast, and in addition to its high sensitivity and precision, CARD-SGS is insensensitive to the frequency of infected cells. Due to the high genetic diversity of HIV in chronic infection, identical RNA molecules can be assumed to originate from the same infected cell. While the CARD-SGS assay provides single-infected cell sensitivity and easily allows interrogation of hundreds of HIV infected cells without *ex vivo* stimulation, there are several important limitations to this study that should be considered. First, the study does not include elite controllers who, with viremia <50 copies HIV RNA/mL, may have additional mechanisms for viral control, including a block to proviral expression of HIV usRNA, that are not seen in VC (7, 41). Second, levels of HIV usRNA were measured by amplifying and quantifying a 1.5kb fragment of *gag-pro-pol* which may be expressed by some defective proviruses as well as by those that are intact. The detection of hypermutated (*i.e.,* multiple G→A mutations, including stop codons) HIV usRNA molecules suggests that at least some defective proviruses can be expressed, as reported by others (17, 21, 42). However, many of the proviruses (about 45-65%) in donors not on ART have been shown to be genetically intact (17-19), in some cases more than 10 years after infection, in stark contrast to the 2-10% of intact proviruses found in individuals on ART (17, 19, 43-46). Another limitation is that our study focused on levels of HIV usRNA in circulating infected PBMC and did not investigate HIV-1 expression in tissues. However, our previous studies of donors on ART showed that the fraction of lymph node mononuclear cells with HIV usRNA is not different than the fraction in PBMC (24). Lastly, we do not know that the HIV usRNA we detect are all LTR-initiated, and not due to expression driven by promoters upstream of the provirus.

In summary, we addressed the question of whether there is a further block to proviral expression in viremic controllers compared to non-controllers by measuring the fraction of infected cells with HIV usRNA. The results of this study reveal that <10% of proviruses express HIV usRNA at a given point in time even in the absence of ART, independent of levels of viremia over a 4,000-fold range, and that viremic control is solely due to having fewer infected cells, likely resulting from a more potent cytotoxic T lymphocyte, natural killer cell, or other innate or adaptive immune response (47-52). These results suggest that at least one mechanism for viremic control varies from that of elite control, in that the additional block to virion production in viremic controllers compared to non-controllers occurs downstream from proviral expression of HIV usRNA, rather than upstream as suggested for elite control (7, 53). Promoting the clearance or elimination of infected cells should be the first priority for achieving HIV control without ART although greater proviral silencing may also be needed to achieve prolonged HIV remission.

## Materials and Methods

### Participant cohort, sample collection, and study approval

PBMC were collected from PWH who were not currently on ART and were enrolled at the University of Pittsburgh in the Optimization of Immunologic and Virologic Assays for HIV Trial (IRB# STUDY20040215) or at the University of California of San Francisco in the SCOPE trial (clinical trial # NCT00187512). The studies were approved by the University of Pittsburgh Institutional Review Board and University of California San Francisco Institutional Review Board. All donors provided written informed consent for HIV quantification and sequencing. All donors were in chronic infection when samples were collection, after the viral setpoint was established. PBMC were separated using Ficoll, resuspended in FBS with 10% DMSO, and stored in liquid nitrogen until testing.

### HIV DNA levels

The number of HIV infected cells was estimated from the levels of HIV DNA in PBMC. HIV DNA levels were measured using the integrase cell-associated single-copy DNA (iCAD) assay (11). Triplicate CCR5 DNA measurements between two dilution factors were performed by BioRad Droplet Digital PCR (54) and averaged to determine the number of cells per aliquot. This step allowed for normalization of HIV DNA measurements to one million PBMC.

### Overview of the cell-associated RNA and DNA single-genome sequencing assay (CARD-SGS)

Although a detailed protocol for CARD-SGS is provided below, this section provides an overview of the method (**Fig. S3**). The CARD-SGS assay was developed as described in Wiegand *et al.* (8) by optimizing methods for the extraction of total nucleic acid from PBMC, efficient degradation of genomic DNA (gDNA) for RNA purification, efficient synthesis of cDNA from cell-associated HIV usRNA using a gene-specific primer in *pol*, and amplification and single-genome sequencing of cell-associated HIV usRNA molecules from multiple aliquots with small numbers of infected cells. Experiments performed to determine the accuracy and sensitivity of the assay for recovery and single-genome sequencing of HIV usRNA are detailed in Wiegand *et al.* (8).

In brief, the CARD-SGS assay is performed by extracting intracellular HIV RNA from ∼8-10 aliquots of one to a few HIV-infected cells with HIV usRNA, followed by single-genome sequencing (SGS) of all HIV usRNA molecules recovered from each aliquot. We perform HIV usRNA SGS on a ∼1.5kb fragment that contains the p6 region of *gag*, protease and the first 700bp of reverse transcriptase in the *pol* gene (p6-PR-RT) as previously described (8). The fraction of cells with HIV usRNA is determined by assuming that identical HIV RNA sequences within an aliquot result from a single-infected cell (due to the high HIV genetic diversity in the donors in untreated chronic infection) and by first estimating the number of total infected cells using the iCAD assay to provide the denominator for number of infected cells in a given aliquot. The levels of HIV usRNA expression in single cells were determined by the number of identical p6-PR-RT sequences in each aliquot. We previously showed that our method can detect a single HIV usRNA molecule in a single cell and that DNA digestion is complete when <300 infected cells are extracted by performing the assay without reverse transcriptase (8). Because the reverse transcription step is known to introduce errors at a rate of about 10^-4^ per sequenced nucleotide of viral cDNA sequences (55), RT-PCR variants that differed by a single nucleotide from a group of 7 or more identical sequences within the same aliquot were considered to belong to the same rake of identical sequences. An overview of the CARD-SGS method is provided in **Fig. S3**. Details of the CARD-SGS protocol as first described by Wiegand *et* al. (8) follow.

### Detailed CARD-SGS method

#### 1. Generating PBMC pellets with few infected cells containing HIV usRNA

Cryopreserved PBMC were thawed by adding warmed RPMI medium dropwise. Thawed cells were serially diluted. The endpoint for HIV usRNA expressing cells was determined empirically by performing RNA extraction, cDNA synthesis, and single-genome amplification on each dilution of PBMC (RNA extraction, cDNA synthesis, and single-genome amplification methods described below). The endpoint was reached when no more than 30% of the PCR reactions in a 96-well plate were positive when the entire cDNA contents were spread across the plate. After the endpoint was determined, multiple aliquots of PBMC at the endpoint were generated, pelleted at 500xg for 5 minutes, supernatant removed, and stored in liquid nitrogen until ready for use.

#### 1.2. RNA extraction

When ready for RNA extraction, the cell pellets stored at an endpoint for HIV usRNA expressing cells were removed from the liquid nitrogen freezer. A working solution of 3M guanidinium hydrochloride (GuHCl+) reagent was prepared from 18.75mL of 8M GuHCL+ (Sigma, #G9284), 2.5mL of 1M Tris-HCl pH 8.0, and 500μL of 100mM CaCl_2_ (Sigma, #21115). A working solution of ∼5.7M guanidinium isothiocyanate (GuSCN+) was prepared from 25mL of 6M GuSCN, 14mL of 1M Tris-HCl pH 8.0, and 53μL of 0.5M EDTA pH 8.0 (AMBION, #AM9260G). For each sample, 100μL GuHCl+ and 5μL 20mg/mL proteinase K (Applied Biosystems, #AM2548) was added to each cell pellet. The samples were vortexed for 10 seconds to resuspend the pellets and incubated at 42°C for 1 hour. Next, 400μL of GuSCN+ and 6μL of 20mg/mL glycogen (Roche, #10901393001) were added to each sample, mixed, and incubated at 42°C for an additional 10 minutes. Nucleic acids were precipitated by adding 500μL of 100% isopropanol and vortexed thoroughly, followed by centrifugation at 21,000x*g* at 21°C for 10 minutes (56). The supernatant was removed, and the pellet was washed with 750μL of 70% ethanol, followed by additional centrifugation at 21,000x*g* at 21°C for 10 minutes. Precipitated nucleic acid pellets were air-dried. Genomic DNA recovery was determined using the Bio-Rad (Hercules, CA) QX200 Droplet Digital PCR System with Bio-Rad 2X ddPCR Supermix for Probes (no dUTP) per the manufacturer’s instructions for CCR5 DNA (57).

For RNA isolation, the nucleic acid pellet was resuspended in 38μL of DNase buffer and 2μL of 10U/μL DNase I (Roche, #04716728001), gently mixed, and incubated for 20 minutes at 37°C. After incubation, 200μL of 6M GuSCN was added to the DNase treated nucleic acid, mixed well, and 250μL of 100% isopropanol was added for precipitation. Samples were vortexed for 10 seconds, then centrifuged at 21,000x*g* at 21°C for 10 minutes to pellet nucleic acid. The supernatant was removed and pellets were washed with 1mL of 70% ethanol, followed by an additional centrifugation at 21,000x*g* at 21°C for 10 minutes. Finally, the RNA pellet was air-dried, then resuspended in 20μL of 5mM Tris-HCl pH 8.0 and used immediately for cDNA synthesis.

#### 1.3. cDNA synthesis

The cDNA was synthesized by the addition of 2.5μL of 10mM dNTPs and 2.5μL of 2μM gene-specific cDNA primer (3500(-) 5’-CTA TTA AGT CTT TTG ATG GGT CAT AA-3’) to each 20μL RNA sample. The RNA was denatured at 85°C for 10 minutes, then immediately cooled at -20°C for 1 minute. Next, 25μL of SuperScript III reverse transcriptase (Invitrogen, #18080-044) master mix was added to the denatured RNA. Master mix was prepared by combining 10μL of 5X first strand buffer, 0.5μL of 0.1M DTT, 13.5μL of RNase-free water, 0.5μL of 40U/μL of RNaseOUT recombinant ribonuclease inhibitor (Invitrogen, #10777-019), and 0.5μL 200 U/μL SuperScript III reverse transcriptase. cDNA was synthesized at 50°C for 1 hour, 85°C for 10 minutes to inactivate the enzyme, cooled to 4°C, then diluted with 150µl of 5mM Tris-HCl pH 8.0 and used immediately or placed at -80°C for temporary storage.

#### 1.4. Single-genome amplification

In a 96-well PCR plate, serial diltutions of cDNA were performed and 10μL PCR reactions were used to determine the endpoint for single molecule amplification. 200µl of endpoint diluted cDNA was then added to a Platinum II Hot-Start Taq Master Mix containing 200μL of 5X Platinum II PCR Buffer, 20μL of 10mM dNTPs, 4μL of each 50μM primer (1849(+), 5’-GGA TCC GAT GAC AGC ATG TCA GGG AG-3’ and 3500(-), 5’-CTA TTA AGT CTT TTG ATG GGT CAT AA-3’), 564μL of molecular-grade water, and 8μL of Platinum II Taq Hot-Start polymerase to amplify single cDNA molecules in a 96-well PCR plate (10μL per well). Thermocycling conditions were as follows: 94°C for 2 minutes, 45 cycles of 94°C for 15 seconds, 60°C for 15 seconds and 68°C for 30 seconds, a final extension of 68°C for 1 minute, then indefinite hold at 4°C. After the first round of PCR, each PCR reaction well was diluted with 50μL of 5mM Tris-HCl pH 8.0. For nested PCR, 2μL of the diluted PCR 1 product was transferred from each well to a new 96-well PCR plate containing 8μL of Platinum II Hot-Start Taq Master Mix (described above) with the nested primers (1870(+), 5’-GAG TTT TGG CTG AAG CAA TGA G-3’ and 3410(-), 5’-CTG TTA GTG GTA TTA CTT CTG TTA GTG CTT-3’). Nested PCR was performed with the following thermocycling conditions: 94°C for 2 minutes, 40 cycles of 94°C for 15 seconds, 60°C for 15 seconds and 68°C for 30 seconds, a final extension of 68°C for 1 minute, then indefinite hold at 4°C. The nested PCR products were diluted with 50μL of 5mM Tris-HCl pH 8.0. Positive PCR products were identified using GelRed (Biotium #41003) detection to illuminate amplified PCR product. Positive PCR reactions were Sanger sequenced as described in Wiegand *et al.* (8).

#### 1.5. Sequence analysis

Sequences were aligned with MAFFT v7.450 (FFT-NS-1 200PAM/k=2 algorithm) in Geneious Prime ® 2020.2.4 (58). Minor adjustments were performed manually. P-distance neighbor joining trees were constructed using MEGA X (59) and rooted on the consensus B HIV sequence. APOBEC3-F/G mediated hypermutation was predicted using the Los Alamos National Laboratory HIV Sequence database tool, Hypermut (https://www.hiv.lanl.gov/content/sequence/HYPERMUT/hypermut.html) (60). Identical sequences obtained from the same cell pellets were inferred to result from a single infected cell (see earlier comment). The number of unique RNA sequences were determined to reflect the number of infected cells with HIV usRNA within each aliquot. The total number of infected cells in each aliquot was previously determined based on the iCAD results described above. The number of infected cells with HIV usRNA was used as the numerator and the total number of infected cells was used as the denominator to determine the fraction of infected cells with HIV usRNA in the samples.

Diversity in plasma RNA, HIV usRNA, and DNA was determined as average pairwise distance (%APD) using an in-house Perl script. Predicted hypermutated sequences were removed and identical sequences were collapsed. HIV-1 DNA sequences were not collapsed prior to APD calculation. To minimize %APD standard deviation at least >10 unique sequences were required for APD analysis (61).

### Single-genome sequencing (SGS) on plasma virus

SGS on plasma virus was performed as described above and in Wiegand *et* al. (8) with the following exceptions: HIV RNA was extracted from a volume of plasma containing ∼5,000 copies of HIV RNA based on the viral loads or from 1mL of plasma in samples with viral loads <5,000 copies/mL. For some individuals, limited plasma sample was available, and few or no plasma RNA sequences were obtained. The rotor for the microcentrifuge was cooled to 4°C prior to use. Plasma was thawed at 37°C and immediately transferred into RNase-free 1.5mL microcentrifuge tubes. Plasma was centrifuged at 5,000xg at 4°C for 10 minutes to remove cellular debris. The supernatant, which contained the viral particles, was removed and placed in new RNase-free 1.5mL microcentrifuge tube being careful to not dislodge the debris. The tubes were centrifuged at 21,000xg at 4°C for 1 hour. The supernatant was removed, and the virion pellets were resuspended in 100μL GuHCl+ and 5μL 20mg/mL Proteinase K (Applied Biosystems, #AM2548). Next, 400μL of GuSCN+ and 6μL of 20mg/mL glycogen (Roche, #10901393001) were added to each sample, mixed, and incubated at 42°C for an additional 10 minutes. Nucleic acids were precipitated by adding 500μL of 100% isopropanol and vortexed thoroughly, followed by centrifugation at 21,000xg at 21°C for 10 minutes (56). The supernatant was removed, and the pellet was washed with 750μL of 70% ethanol, followed by additional centrifugation at 21,000xg at 21°C for 10 minutes. Finally, the RNA pellet was air-dried, then resuspended in 20μL of 5mM Tris-HCl pH 8.0 and used immediately for cDNA synthesis (following **Materials and Methods** Section 3: cDNA synthesis). The entire viral RNA extraction was used for cDNA synthesis as described above. The cDNA was serially diluted to an endpoint (<30%) on 96-well PCR plates and amplified (following **Materials and Methods** Section 4: single-genome amplification) and the products were Sanger sequenced as in Wiegand *et al.* (8).

### Sequence data availability

All sequence data are available on GenBank at accession numbers: XXXX-XXXX

*Defining the cut-off for the number of HIV usRNA in “high-expressing” cells as >10 copies* In order to define the cut-off for the number of HIV usRNA in “high-expressing” cells, the distribution of HIV usRNA copies per cell in NCs was used. The count distribution was Poisson-like. Cells with >10 RNA molecules per cell were statistically more rare than cells with <10 RNA molecules (Bonferroni-corrected p-value of 0.034). Therefore, cells that produced >10 HIV usRNA copies per cell were defined as “high-expressing”.

## Statistical analyses

The Mann-Whitney U-test, Wilcoxon signed rank test, chi-square test, and Spearman’s correlation were performed using Prism GraphPad 9.2. Spearman’s correlation was selected due to the random variation in biological systems. Statistical tests applied are indicated in the text and/or figures/table legends.

To quantify the number of HIV usRNA molecules in each aliquot of infected cells assayed, the sequence alignment containing HIV usRNA undergoes “defuzzing” (adding the ‘fuzz’ sequences into the identical sequence rake) where the number of unique HIV usRNA molecules is determined per aliquot using the R script available at https://github.com/michaelbale/RStuff/blob/master/fPECS_fncs.R. It is then possible to calculate the fraction of provirus expressing cells by taking the ratio of unique HIV usRNA sequences in the same aliquot as the numerator and number of HIV DNA molecules in each respective aliquot as the denominator. Additional statistical analyses were performed using R.

## Acknowledgements

We would like to thank the donors for participating in this study. We would also like to thank Connie Kinna, Valerie Turnquist, Ann Wiegand, and Susan Toms for administrative support, Beth Fyne for providing valuable reagents and cell lines, Laura Newman and Leslie Lipkey for sequencing, and Valerie Boltz, Jason Rausch, and Frank Maldarelli for consultation.

Figure S3 was modified from Wiegand *et al.* (8) and created by using resources from biorender.com.

## Author Contributions

AAC, AW, FH, JMC, JWM, MFK designed the study. AAC, AW, FH, JJ, JS, AW, AC, RG, and EKH performed experiments. AAC, AW, FH, BTL, RG, JMC, JWM, and MFK analyzed results. AAC, AW, MJB, and WS performed sequence data analyses. MDS, RH, EKH, SGD, and JWM provided patient samples. AAC and MFK wrote the manuscript. All authors reviewed the manuscript.

## Classification

Biological Sciences, Microbiology.

---

**Supplemental Figure 1.**
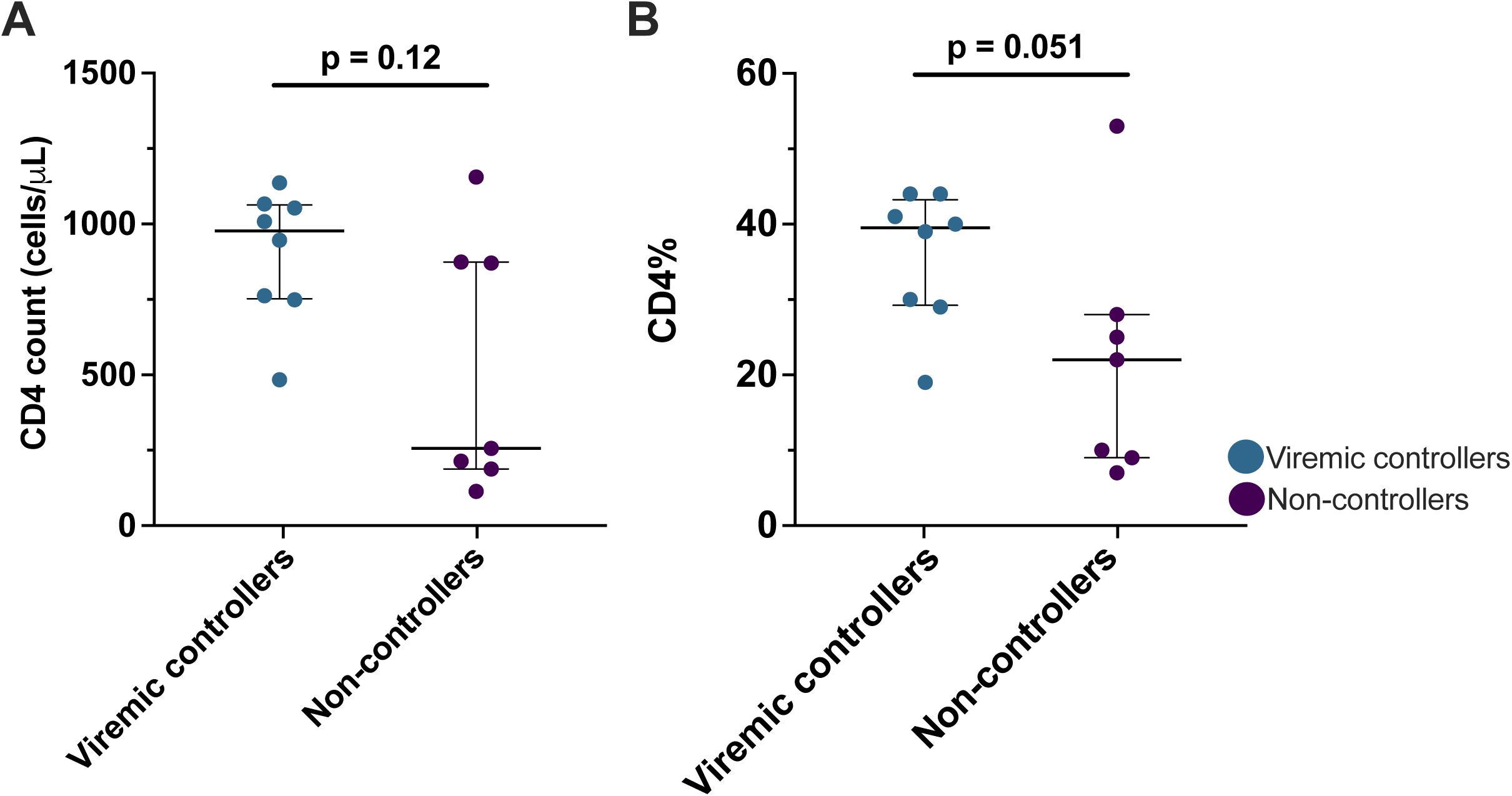
M*easurement of CD4+ T cells in untreated donors.* Clinical assays using flow cytometry measured the (**A**) CD4+ T cell count as cells/μL, and (**B**) %CD4+ T cells in PBMC in whole blood as measured by flow cytometry. Mann-Whitney test was performed with the median and interquartile range reported. Each symbol represents an individual donor within the respective group. Measurements are at the time of sampling for **Table 2**.

**Supplemental Figure 2.**
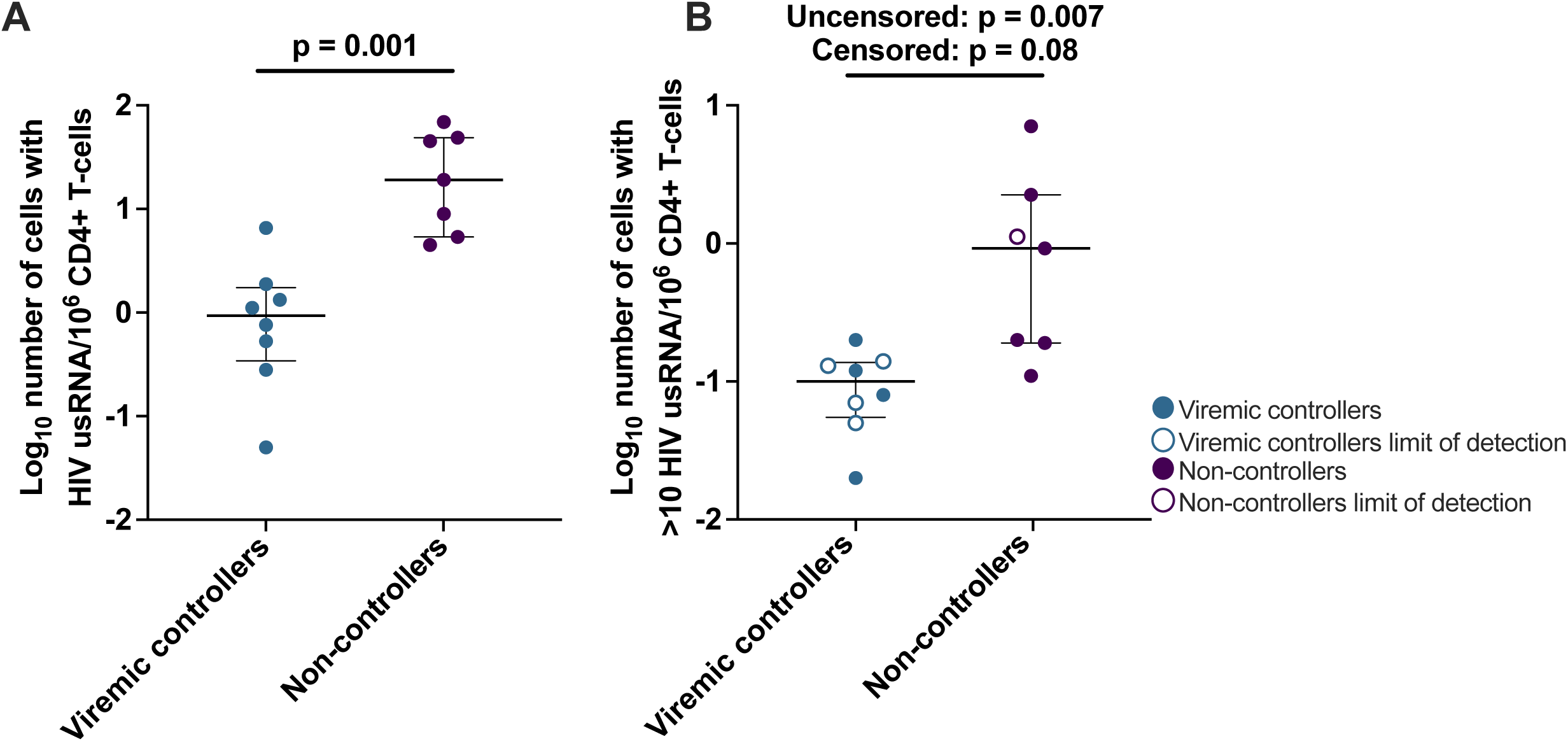
N*umber of cells with HIV usRNA normalized to CD4+ T cells.* **(A)** The number of cells with HIV usRNA per million PBMC was normalized to CD4+ T cells by the %CD4 at the time of sampling. **(B)** The number of cells per million PBMC that are “high-expressing” (>10 copies HIV usRNA) normalized to CD4+ T cells by the %CD4 at the time of sampling. Open shapes indicate no detectable “high-expressing” cells in a particular donor, plotted at the estimated limit of detection estimate determined as 1/number of PBMC assayed. Mann-Whitney test for both uncensored and censored data (donors with undetectable “high-expressing” cells, open shapes) with median and interquartile range as shown. Each symbol represents an individual donor within the respective group.

**Supplemental Figure 3.**
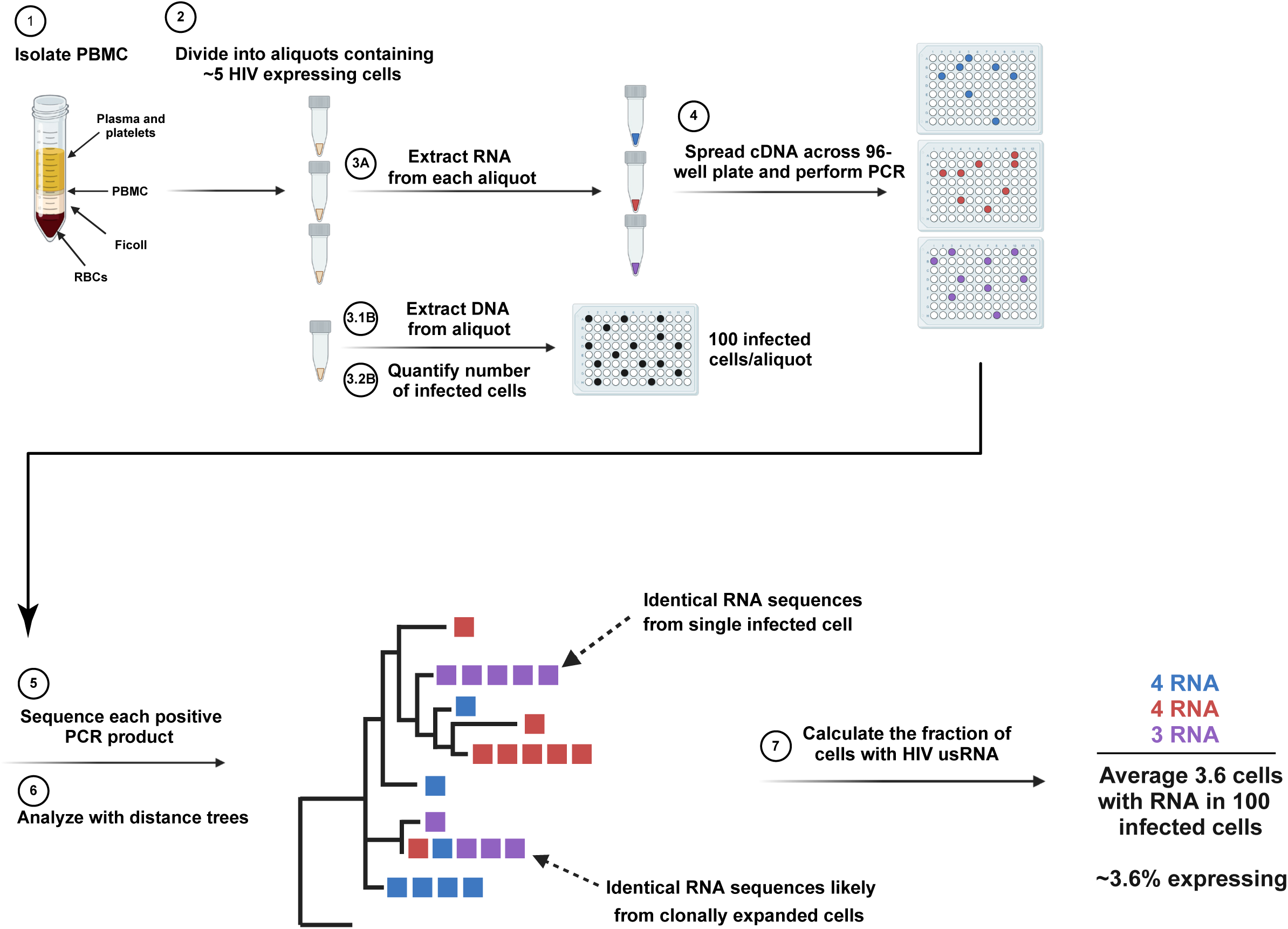
F**low chart of CARD-SGS assay.** PBMC are isolated from whole blood and separated into separate aliquots. One aliquot is used to determine the number of infected cells per aliquot. Then separate aliquots are made with ∼5% HIV usRNA expressing cells per aliquot. RNA is extracted, treated with DNAse, and undergo cDNA with a gene-specific primer. The synthesized cDNA is spread across a 96-well plate and PCR amplified. Following two rounds of PCR targeting the p6-PR-RT region, amplicon detection of positive wells are determined. PCR positive wells are selected and undergo Sanger sequencing. Electropherograms are analyzed through custom in-house scripts, alignment is generated, with neighbor-joining distance trees reconstructed. Analysis of the tress distin-guishes identical HIV usRNA sequence from the same aliquot, indicating they derived from a single infected cell; or from different aliquots suggesting they were derived from clonally expanded cells. The sum of the number of unique HIV usRNA populations are divided over the number of infected cells analyzed to determine the fraction of cells expressing HIV usRNA at the time of sampling.

